# Characterization of immunity-inducing rhizobacteria highlights diversity in plant-microbe interactions

**DOI:** 10.1101/2024.05.23.595641

**Authors:** Mackenzie Eli William Loranger, Winfield Yim, Matthew Toffoli, Marie-Christine Groleau, Arvin Nickzad, Nadia Morales-Lizcano, Thomas Berleth, Wolfgang Moeder, Eric Déziel, Keiko Yoshioka

**Author notes:** Co-first authors (Mackenzie Eli William Loranger, Winfield Yim). Additional Email address.

## Abstract

The narrow region of soil surrounding roots (rhizosphere) contains an astonishing diversity of microorganisms. Some rhizosphere bacteria can improve plant health and immunity, via direct competition with pathogens or by establishing heightened immunity in aboveground tissues, a phenomenon known as Induced Systemic Resistance (ISR). We screened a bacterial library from agricultural soils to identify strains that, after root treatment, induce immunity in *Solanum lycopersicum* (tomato) against the fungal pathogen *Botrytis cinerea*. Here, we report the establishment of a screening method and characterization of a subset of five strains, belonging to the species *Bacillus velezensis, Paenibacillus peoriae* and *Pseudomonas parafulva*, that induced systemic resistance in tomato. However interestingly, only two of them triggered canonical ISR in Arabidopsis, indicating plant host specificity and/or alternative modes of actions. Furthermore, some of the strains displayed direct anti-microbial activity. We also found the requirement of the lipid-binding protein DIR1 in ISR establishment, indicating a possible convergence of SAR and ISR signaling. Finally, we found that *P. parafulva* TP18m, also displayed strong effects on root development. Taken together, we have identified taxonomically diverse immunity-inducing bacteria. Our characterization revealed diverse features, highlighting the complexity of bacteria-host interaction in the rhizosphere.

**Highlight:** We identified taxonomically diverse rhizobacteria that induce systemic resistance in tomato plants to Botrytis after application to the root. These bacteria display diverse modes of action to improve plant health.

## Introduction

The rhizosphere, the narrow region of soil surrounding plant roots, contains an astonishing number and variety of microorganisms, referred to as the plant microbiota; their total pool of genetic material is referred to as the microbiome (Berendsen et al., 2012). Plant root secretions influence the composition of the microbiota and consequently the rhizosphere microbiota differs significantly from the surrounding soil (Pieterse et al., 2014). In turn, the microbiota significantly affects plant growth, development, nutrient- and water-acquisition, as well as tolerance to abiotic and biotic stresses such as drought, temperature, and pathogens (Trivedi et al., 2020). The rhizosphere microbiome provides crucial protection to plants from pathogenic organisms in two main ways. First, these microbes work as a front line of immunity against soil-borne pathogens through direct competition among microorganisms. Second, the rhizosphere microbiome can elicit natural immunity in plants (Pieterse et al., 2014; Trivedi et al., 2020).

Like animals, plants have evolved elaborate defense systems against pathogens. Plants possess a complex system that directly or indirectly recognizes numerous microorganisms. A first line of defense is via the recognition of common microbial compounds, such as bacterial flagellin or fungal chitin, called pathogen- or microbe-associated molecular patterns (PAMPs or MAMPs), via pattern recognition receptors (PRR), which induce PAMP-triggered immunity (PTI). For example, the PRR FLS2 recognizes a 22 amino acid epitope of the bacterial flagellin (flg22) (DeFalco and Zipfel, 2021). However, many pathogens have evolved effector proteins that, when secreted into the plant cell, can dampen the plant immune system; in turn, plants have acquired a second layer of defense, where NB-LRR (nucleotide binding-leucine rich repeat) receptor proteins can recognize these effectors and trigger a stronger type of immunity termed effector induced immunity (ETI) (Ngou et al., 2022). Systemic acquired resistance (SAR) and induced systemic resistance (ISR) are two types of induced systemic resistance against pathogens. Once plants establish SAR or ISR, they react faster and more strongly to pathogen infection (Vlot et al., 2021), a phenomenon often referred to as “priming” (Conrath et al., 2006). SAR is induced by prior infection with avirulent strains of pathogenic microorganisms that trigger ETI and is effective mainly against biotrophic pathogens (pathogens that require living host cells) via the defense phytohormone salicylic acid (SA)(Vlot et al., 2021). By contrast, ISR is induced by non-pathogenic microorganisms closely associated with root systems and confers broad-spectrum resistance against biotrophic and necrotrophic pathogens (pathogens that do not require living host cells, and thus actively kill host cells and derive energy from the dead cells), as well as against herbivores, at the site of induction and in systemic tissues (Haney et al., 2018; Pieterse et al., 2014).

Since the discovery of ISR in 1991, ongoing research has elucidated some of the mechanisms of its activation, and identified various elicitors, such as siderophores, biosurfactants, volatiles, and antibiotics (Pieterse et al., 2014). The best characterized ISR-inducing bacterium is *Pseudomonas simiae* WCS417, which was originally isolated from wheat roots (Pieterse et al., 2021). WCS417 - like many rhizobacteria - possesses both ISR-inducing as well as plant growth promoting (PGP) activities (Pieterse et al., 2021). Plants grown with WCS417 exhibit larger biomass and increased lateral root formation (Pieterse et al., 2021). PGP Rhizobacteria (PGPRs) frequently produce phytohormones including auxins, gibberellins, and cytokinins. Via ACC deaminase, PGPRs can also reduce ethylene production in roots. All of these affect the plant root development. Furthermore, PGPRs assist the host plant in nutrient uptake by fixing nitrogen and solubilizing or chelating minerals, such as phosphate (P_i_), potassium and iron via the production of organic acids and siderophores (Backer et al., 2018).

The signal transduction of ISR is still not well understood. Initial studies showed that ISR - unlike SAR - is dependent on jasmonic acid (JA) and ethylene (ET) signaling, rather than salicylic acid (Pieterse et al., 1998). Interestingly, *NON EXPRESSOR OF PR GENES1* (*NPR1*), the master regulator of SA responses, is also involved in ISR (Pieterse et al., 1998). Further studies revealed that the JA-associated transcription factor *MYC2* is a central regulator (Pozo et al., 2008) in systemic leaves. Another transcription factor, *MYB72,* is primarily induced in the roots and is essential for systemic signaling (Van Der Ent et al., 2008; Zamioudis et al., 2015).

In current agricultural systems, diseases claim ∼20–40% of crop yields (Savary et al., 2019), and climate change may exacerbate these losses by increasing the spread of pathogens and subjecting plants to increased abiotic stresses, thereby decreasing their ability to fight pathogens (biotic stress). The fungal necrotrophic pathogen *Botrytis cinerea,* commonly known as gray mold, is a major concern for agriculture, causing widespread crop loss in a broad range of species and therefore has a destructive economic impact (Fillinger and Elad, 2016).

Over the last decades, chemical pesticides have been extensively used to protect crops from pathogens, but they also have detrimental effects on the environment and human health (Tudi et al., 2021). Thus, one approach to enhance pathogen resistance is by priming the plant’s own immunity. While SAR often causes a fitness penalty, ISR provides a natural immunity without any significant cost to the plant that can be used for crop protection from a wide range of diseases (Berendsen et al., 2012).

We set up a pipeline to screen bacteria from a library of culturable bacteria from various environments in Canada for their ability to induce systemic resistance against *B. cinerae* in *Solanum lycopersicum* (tomato); and here we report a detailed characterization of five strains that had previously been shown to display antagonistic activity against phytopathogenic *Xanthomonas* species (Olishevska et al., 2023).

## Material & Methods

### Plants and growth conditions

*Solanum lycopersicum* (tomato) cv. ‘Glamour’ seeds were germinated in sand at 22°C with a 16- hour light cycle. Two-week-old seedlings were transplanted to soil or Jiffy pellets.

Arabidopsis plants were grown in a growth chamber at 22°C with an 8-hour light cycle. *Arabidopsis thaliana* Col-0, *jar1* (Staswick et al., 1992) *ein2* (Alonso et al., 1999) *sid2* (Wildermuth et al., 2001), *rbohd* (Angel Torres et al., 2002.), and *dir1-2* (Maldonado et al., 2002) were used in this study.

### Bacterial treatments and Botrytis infection

Bacterial isolates were cultures on Lysogeny Broth (LB) agar at 28°C. The bacteria were resuspended in 10 mM MgSO_4_ at an OD_600_ of 0.1. Peat pellets (Jiffy-7, 42 mm pots) were soaked in 50 mL of this inoculum before transplanting 2-week-old tomato seedlings. Eight biological repeats were performed for each treatment with 1 plant per peat pellet. One week after the bacterial inoculation, the peat pellets with the plants were transplanted into larger pots with soil and maintained for another 2 weeks until ready for the challenge infection with *B. cinerea.* As controls, plants were mock inoculated with sterile 10 mM MgSO_4_ solution or with the positive ISR control strains, *P. defensor* WCS374 (Berendsen et al., 2015).

*Botrytis cinerea* isolate MEE B191, was used for this study (provided by Canadian Collection of Fungal Cultures, Agriculture and Agri-Food Canada, Ottawa, ON, Canada). *B. cinerea* was grown on potato dextrose agar (PDA) for seven days at 24°C. To prepare the inoculum, fungal conidia were collected in Sabouraud maltose broth (SMB) by scraping off fully grown plates and adjusted to a final concentration of 2.5x10^5^ conidia mL^-1^. Four to five-week-old tomato plants were challenge inoculated by placing a 10 μl droplet of conidia onto the midvein of detached adaxial leaflets. The inoculated leaves were kept in sealed Petri dishes lined with sterile water-saturated filter paper to maintain a high humidity environment. The Petri dishes were kept in 24h light conditions at room temperature for 3-4 days. Then, photos of the Petri dishes were taken, and lesion size was analyzed using the Colour-analyzer tool (Loranger et al., 2024). For each treatment, two leaves each of eight plants were used resulting in 16 biological repeats. 4-week-old Arabidopsis were soil drenched with 50 mL of bacterial suspension adjusted to an OD_600_ of 0.2. After one week incubation, the *B. cinerea* challenge inoculation was performed on attached leaves by adding a 10 μl droplet of inoculum to three leaves of four individual plants. The flat that contained the plants was then domed and sealed to maintain high humidity. The inoculum was prepared as described above; however, the final concentration was increased to 2.5x10^6^ conidia mL^-1^. The plants were kept in 24h light conditions at room temperature for 3-4 days. Images were taken and the width and heigh of the circular lesions were measured in Fiji to calculate the approximate area of the lesion (Schindelin et al., 2012).

### PGP assays

Seeds were germinated on MS agar plates, and 3-5-day-old germinated seedlings were transplanted to new MS agar plates. Bacteria were resuspended in 10 mM MgSO_4_ at an OD_600_ of 0.1 and 15 μL droplets of inoculum were deposited 3 cm from the root tip. Plates were than placed upright in a growth rack under constant light at 24°C. Root measurements were performed 14 days after the treatment.

### Direct antagonism assays

When testing anti-fungal activity against *B. cinerea,* the bacterial strains were resuspended in 10 mM MgSO_4_ and diluted to a final OD_600_ of 0.1. In a 9.5 cm diameter Petri dish with PDA agar, a 4 mm wide plug of *B. cinerea* was placed into the center of the plate. Then 2.5 cm away from either side a streak of bacteria was added, effectively flanking the fungal plug on both sides.

These were incubated for 6 days before photographs were taken.

A lawn of *P. syringae* was plated onto LB plates before placing a droplet (OD_600_ = 0.8) of each bacterial strain into the center. The plates were incubated at 28°C overnight before photographs were taken.

### GUS histochemical staining

The *proMyb72:GUS* line described in Zamioudis et al., (2015) was used. GUS histochemical staining was done using a staining solution composed of 50 mM sodium phosphate buffer (pH 7.0), 2 mM potassium ferrocyanide, 2 mM potassium ferricyanide and 2 mM X-Gluc (modified from Kim et al. (2006). Samples were incubated at 37°C for 3 h. Stained roots were stored in 70% ethanol and imaged using the Leica MZ16F stereomicroscope.

### Phosphate solubilization

The bacterial isolates were screened for their calcium phosphate solubilizing activity on Pikovskaya medium (PVK). Isolates were aseptically spot inoculated three times on each plate, approximately equidistant from each other. All the plates were incubated at 28°C for 4 days (modified from Nautiyal (1999)).

### Siderophore production

Siderophore production was determined by use of the Chrome Azurol S (CAS) assay. On CAS agar, siderophores remove iron from the CAS dye complex, resulting in a blue-to-orange color change in zones surrounding the colonies (Vial et al., 2008). Strains were grown overnight in TSB at 30°C. Cells were washed twice in 0.8% NaCl and suspensions were adjusted to an OD_600_ of 2.0 A 3 μl droplet was spotted onto CAS agar. The plates were incubated for 24 h at 30°C. The production of siderophores was estimated by measuring the area of the halo (mm^2^) surrounding the colonies. Some strains were unable to grow on CAS agar. For those, a modified method was used as described in (Milagres et al., 1999).

### ACC deaminase and Nitrogen Fixation Assays

ACC deaminase production was determined on DF agar as described with either ACC or (NH_4_)_2_SO_4_ as nitrogen sources (Gupta and Pandey, 2019). Nitrogen fixation was performed in the same conditions but without addition of a nitrogen source. Strains were grown overnight in TSB at 30°C. Cells were washed twice in 0.8% NaCl and suspensions were adjusted to an OD_600_ of 2.0. A 3 μl droplet was spotted on each media. Plates were incubated at 30°C and monitored each day for a week.

### Primers

Strain specific primers are listed in Table S1

### Statistical Analysis

To evaluate the effects of each bacterial strain on its ability to enhance resistance against *B. cinerea* in tomato and Arabidopsis, each treatment was compared to the buffer-inoculated control plants. Differences between treatments and buffer-treated plants were evaluated using a one-way ANOVA followed by a Dunnet’s test post-hoc. Statistical significance was fixed at a probability level of p<0.05. To evaluate the PGP effects through the comparison of primary root length and lateral root number a one-way ANOVA followed by a Tukey’s HSD test post-hoc. All statistical analyses were performed in R (v. 4.0.2).

## Results

The use of rhizobacteria for crop protection has recently gained a lot of attention; however, the effectiveness of any biological control agents strongly depends on climate, location, and type of crop (Lahlali et al., 2022). Tomatoes are an important vegetable crop in Canada. Therefore, we used a library of bacteria derived from Canadian agricultural soil samples (Olishevska et al., 2023) to identify strains that can induce systemic resistance in plants after application to the root. Fig. 1A summarizes our established workflow: 1. Inoculation - Transfer two-week-old tomato (*Solanum lycopersicum* cv. ‘Glamour’) seedlings to compressed peat pellets (Jiffy-pellets, Jiffy Growing Solutions, Zwijndrecht, the Netherlands) that had been soaked in 50 mL/pellet of a bacterial inoculum. 2. Colonization - one week later these pellets were transplanted to larger pots with soil, and 3. Pathogen inoculation - 1-2 weeks after colonization, two leaves per plant were collected from 8 plants per treatment. The leaves were then challenge-inoculated with a solution of conidia from *B. cinerea* MEE B191. Droplets (10 μl) of the conidia solution were placed on the midvein of detached leaves. The inoculated leaves were placed on water-saturated filter papers and sealed in Petri dishes to maintain high humidity. The lesion development on the detached leaves were analyzed after 3-4 days using our Colour-analyzer pipeline (Loranger et al., 2024). Colour-analyzer is particularly useful for asymmetric lesions and high through put analyses. Briefly, the Petri dishes with the leaves were photographed, and these images were analyzed in the Colour-analyzer tool (https://vittorioaccomazzi.github.io/LeafSize/). Colour- analyzer uses two colour models, utilizing both HSV (*Hue, Saturation, Value*) and L*a*b* values. The *a* b\** values were used to distinguish between infected and healthy tissue, while the *H* and *S* channels were used to differentiate the leaf area from the background. The tool analyzes each leaf individually, producing a CSV file in which each leaf will have both the lesion area and total area indicated (Loranger et al., 2024).

**Figure 1.**
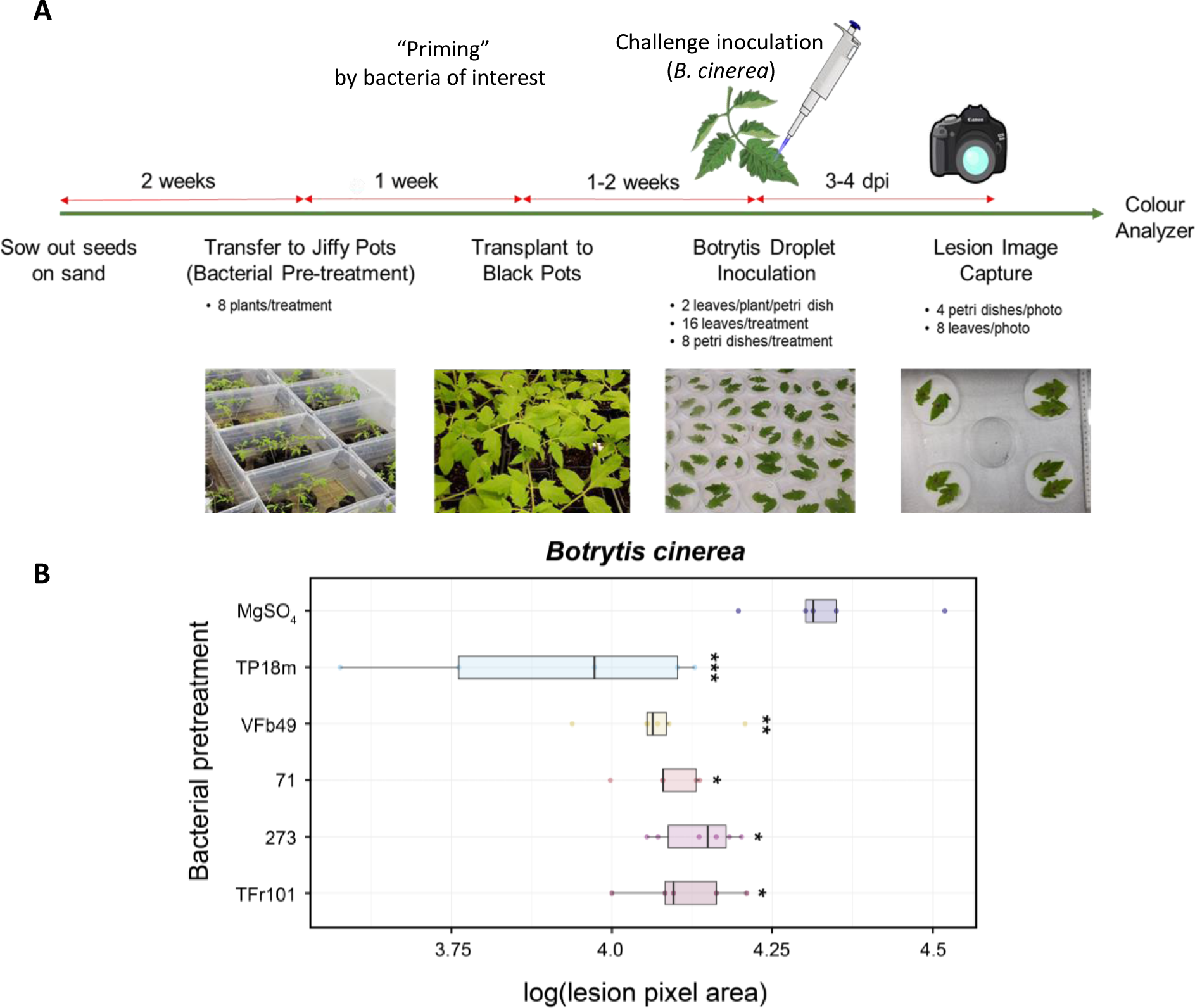
Screening of bacterial isolates that can induce systemic resistance in tomato. **A.** Schematic of the screening methodology in tomato. Two-week-old seedlings were transplanted into Jiffy pellets that had been drenched in a suspension of bacteria (OD^600^ = 0.1). After one week the plants were transplanted to pots. One to two later, leaflets were transferred into Petri dishes containing a wet filter paper. A drop of 10 μl of a solution of *B. cinerea* conidia (2.5x10^5^ mL^-1^) was placed onto the midvein of leaflets. After 3-4 days of incubation in sealed Petri dishes, photos were taken, and the lesion size was analyzed using the Colour-analyzer tool. **B.** Boxplot representing lesion area induced by *B. cinerea*. (n = 6; p < 0.001 ***, < 0.01 **, < 0.05 *. ANOVA followed by post hoc Dunnett’s test)

So far, about 500 bacterial strains have been tested and here we report on a subset of five strains that had previously been shown to display antagonistic activity against several Xanthomonas strains (Olishevska et al., 2023). They induced a statistically significant reduction in *Botrytis* lesion size in at least three independent experiments (Fig. 1B). These were two *Bacillus velezensis* strains (71 and VFb49), two strains of *Paenibacillus peoriae* (273 and TFr101), and *Pseudomonas parafulva* TP18m.

*Pseudomonas defensor* WCS374, which was originally isolated from the rhizosphere of potato plants, is one of the best studied ISR inducers (Berendsen et al., 2015). We compared one of our strong immunity-inducing strains, *B. velezensis* VFb49, with WCS374 and found the same degree of lesion reduction in tomato cotyledons infected with *B. cinerea,* while *E. coli* treatment (negative control) did not show any effect (Fig. 2). This indicates that *B. velezensis* VFb49 has strong biocontrol capacity.

**Figure 2.**
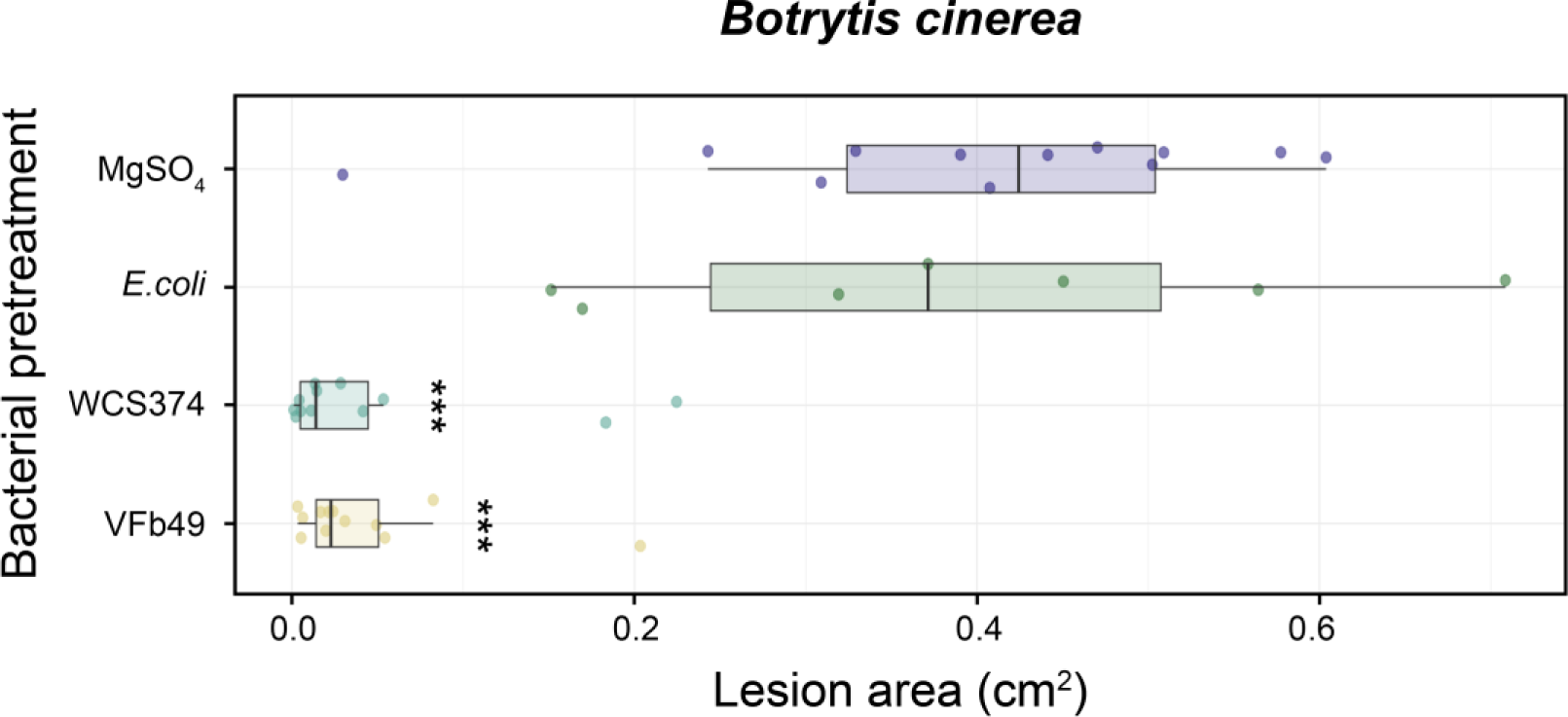
Induction of systemic resistance in tomato seedlings after *Bacillus velezensis* (VFb49) and *Pseudomonas defensor* WCS374 treatment. One-week-old tomato seedlings were transplanted into Jiffy pellets that had been drenched in a solution of bacteria (OD_600_ = 0.1). After one week, a drop of 10 μl of a solution of *B. cinerea* conidia (2.5^5^x10 mL^-1^) were placed onto the midvein of intact leaflets. The plants were domed and sealed, after 3-4 days, the lesion area was measured. Boxplot representing lesion area induced by *B. cinerea*. (n = 6; p < 0.001 ***, < 0.01 **, < 0.05 *. ANOVA followed by post hoc Dunnett’s test)

### Direct antimicrobial activity

Since these strains had previously been shown to possess anti-*Xanthomonas* activity (Olishevska et al., 2023), they were also assessed for direct antimicrobial activity against *B. cinerea*. A plug of fungal mycelium was placed in the center of a Petri dish and bacteria were streaked out on both sides of the plug. After 6 days the growth of the *B. cinerea* mycelium was assessed (Fig. 3A). In this assay, *B. velezensis* strains 71 and VFb49 displayed very strong inhibition of fungal growth, while *P. peoriae* strains 273 and TFr101 had a less pronounced effect, and *P. parafulva* TP18m did not exhibit antimicrobial activity against *B. cinerea*. A known ISR-inducing strain, *Pseudomonas simiae* WCS417 (Pieterse et al., 1996; Pieterse et al., 2012) also displayed an intermediate level of inhibition, while *E. coli* did not affect fungal growth (Fig. 3A). We further assessed the antimicrobial activity of these strains against the bacterial tomato pathogen, *Pseudomonas syringae* pv. tomato DC3000. A droplet of bacterial solution was added to a lawn of *P. syringae* and the formation of a halo around the droplet was assessed. Only the two *Paenibacillus* strains 273 and TFr101 displayed anti-*Pseudomonas* activity in this assay (Fig. 3B).

**Figure 3:**
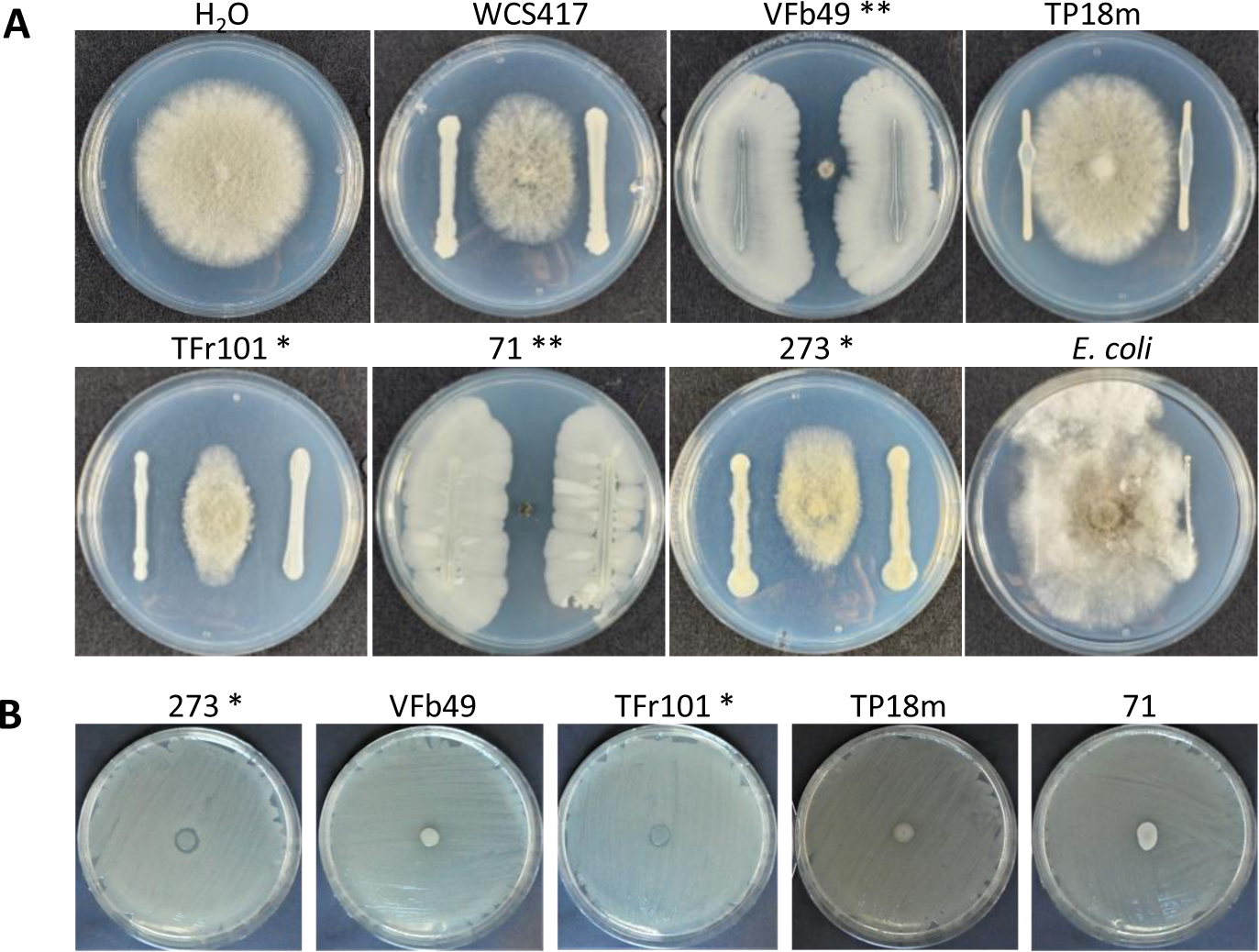
Direct antimicrobial activity of ISR-inducing strains against *B. cinerea and Pseudomonas syringae* pv. tomato DC3000. **A.** Direct competition between bacterial strains and *B. cinerea*. In a Petri dish with PDA agar, a plug of *B. cinerea* was placed into the center of the plate and on either side a streak of bacteria was added. Photographs were taken after 6 days. Experiments were conducted with 3-5 plates per strain. Similar results were obtained at least three times. ** denotes strong inhibitory effect on *B. cinerea* growth. * denotes weaker but consistent inhibition. *P. simiae* WCS417 and *E. coli* served as controls. **B.** Direct antagonism assay against *Pseudomonas syringae* pv. tomato. A lawn of *P. syringae* was plated onto LB plates before placing a droplet of each bacterial strain into the center. The plates were incubated at 28°C overnight before photographs were taken.

### Effect in Arabidopsis and factors required to induce immunity

Some ISR-inducing strains are effective across a range of plant hosts, while others are limited to one or few host plants (Afzal et al., 2019; Berendsen et al., 2015). Furthermore, there is limited information regarding the molecular mechanisms of ISR signaling in tomato. Thus, to dissect the mechanism and signal transduction of resistance induced by our strains, we used the well-studied model plant, *Arabidopsis thaliana*. Interestingly, only *B. velezensis* VFb49 and *P. parafulva* TP18m induced ISR in Arabidopsis accession Columbia wild type (Col-0 WT), as did the known ISR inducer *P. simiae* WCS417 (Fig. 4).

**Figure 4.**
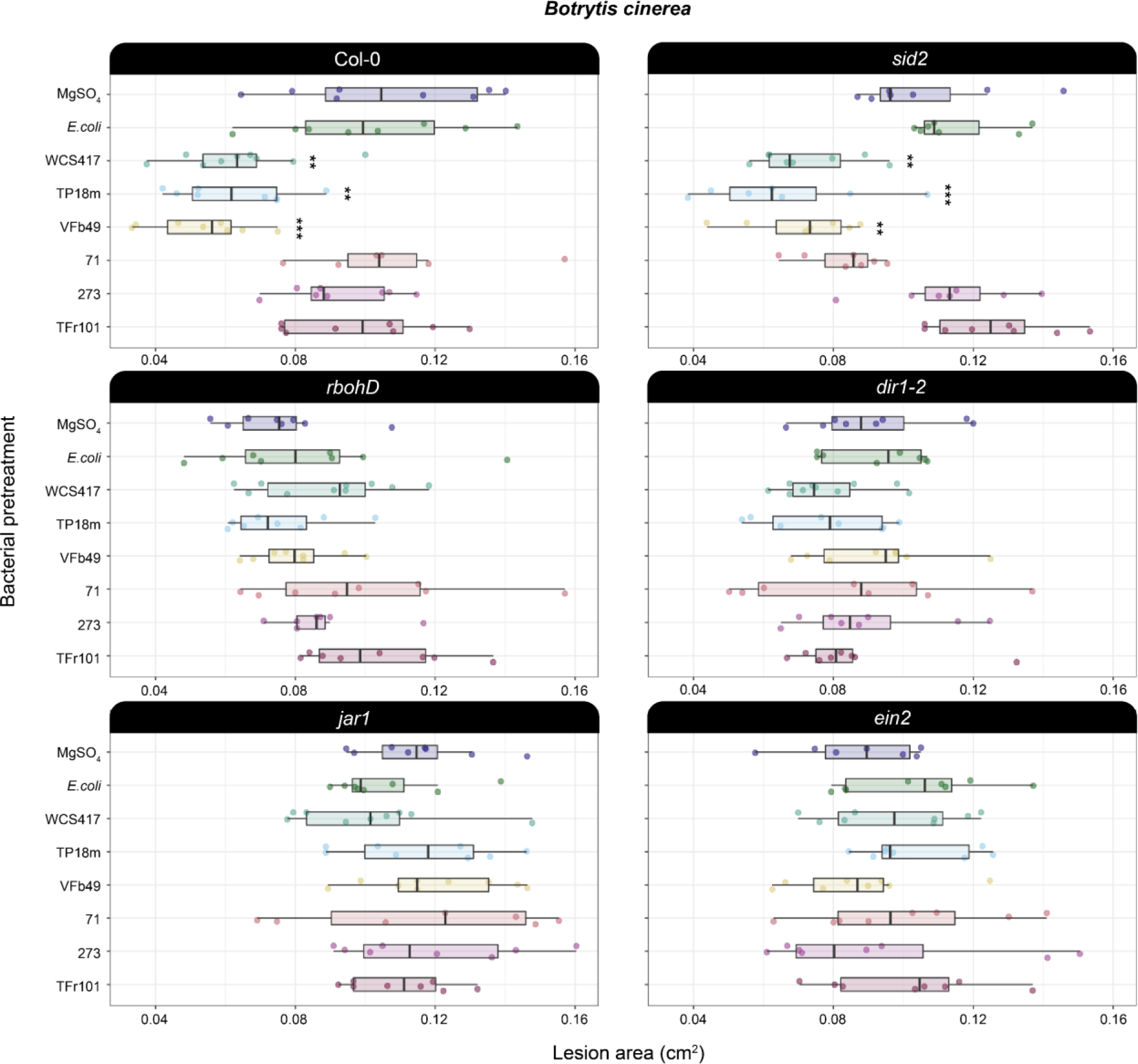
Boxplot depicting *B. cinerea* lesion development after bacterial pre-treatment in Arabidopsis signaling mutants. 4-week-old Arabidopsis were soil drenched with bacterial suspension (OD_600_ = 0.2). After one week a 10 μl droplet of *B. cinerea* conidia (2.5x10^6^ conidia mL^-1^.) was placed on attached leaves of Arabidopsis plants. The plants were domed and kept in 24h light conditions at room temperature, after 3-4 days, the lesion area was measured. (Col-0 - Wt, *sid2* - SA biosynthesis, *rbohD* – NADPH oxidase (ROS), *dir1-2* - *defective* in *induced resistance1*, *jar1* – JA signaling deficient, *ein2* – ET signaling mutant) . Difference between MgSO_4_ and each bacterial treatment was evaluated within genotypes using a one-way ANOVA followed by a post hoc Dunnett’s test. (n = 8-12; p < 0.001 ***, < 0.01 **, < 0.05 *).

ISR has been studied extensively in Arabidopsis (Hacquard et al., 2017; Pieterse et al., 2014). It is believed to be regulated by JA/ET signaling and is mostly independent of SA (Pieterse et al., 2012). However, most of this research was based on a few *Pseudomonas fluorescens* strains (Berendsen et al., 2015), and recent data using other bacterial species suggest that it is possible that depending on the inducing bacterial strains, different phytohormones and their signaling pathways may be required (Nie et al., 2017; Niu et al., 2011; Vlot et al., 2021). Therefore, we tested the efficacy of our strains in various Arabidopsis mutants related to JA, ET and SA signaling, as well as other immunity-related genes. As for *P. simiae* WCS417, ISR was abolished for our two ISR-positive strains in the *jar1* (JA insensitive) and *ein2* (ethylene insensitive) mutants, while no effect was observed in the SA biosynthesis mutant *sid2* (Fig. 4). Thus, *B. velezensis* VFb49 and *P. parafulva* TP18m activate ISR in Arabidopsis through the same or a similar pathway as *P. simiae* WCS417.

Recently, the importance of reactive oxygen species (ROS) in root colonization of beneficial bacteria has been reported (Tzipilevich et al., 2021); thus, we also tested a mutant for RBOHD, which encodes a plasma membrane localized NADPH oxidase that produces a burst of reactive oxygen species (ROS), which plays a crucial role in the induction of PTI (Angel Torres et al., 2002). Interestingly, all three strains failed to induce ISR in the *rbohD* mutant (Fig. 4), suggesting that ROS production also plays an important role in ISR induction in a range of bacterial species.

In addition, a connection of the SAR-related lipid transfer proteins EARLY ARABIDOPSIS ALUMINUM INDUCED 1 (EARLI1) and AZELAIC ACID INDUCED 1 (AZI1) with ISR had recently been suggested (Cecchini et al., 2015; Vlot et al., 2021); thus, we also tested the SAR-deficient *dir1* (*defective* in *induced resistance1*) mutant (Maldonado et al., 2002). *DIR1* encodes an apoplastic lipid transfer protein and is believed to be part of the systemic signal to establish SAR (Vlot et al., 2021), however its role in ISR has not been tested. Interestingly, all three strains did not trigger ISR in *dir1*, suggesting that DIR1 plays a role in ISR, and that there is some overlap between SAR and ISR signaling.

During ISR establishment very few transcriptional changes are known; however, the *MYB72* transcription factor is induced transcriptionally, and is essential for ISR signaling. It is specifically activated in the roots upon colonization by strain WCS417 (Van Der Ent et al., 2008). Thus, we tested whether *MYB72* expression is induced by our strains using a *pMYB72*:*GFP*lJ*GUS* reporter line (Zamioudis et al., 2015). Corroborating the ISR induction against *B. cinerea* (Fig. 4), only *P. simiae* WCS417*, B. velezensis* VFb49 and *P. parafulva*

TP18m induced the GUS reporter gene in the root 3 days after bacterial treatment (Fig. 5). Thus, two strains, *B. velezensis* VFb49 and *P. parafulva* TP18m, act though the canonical ISR pathway similarly to *P. simiae* WCS417 (Pieterse et al., 2021), while the three other strains are either specific for tomato plants or at least do not trigger ISR in Arabidopsis.

**Figure 5.**
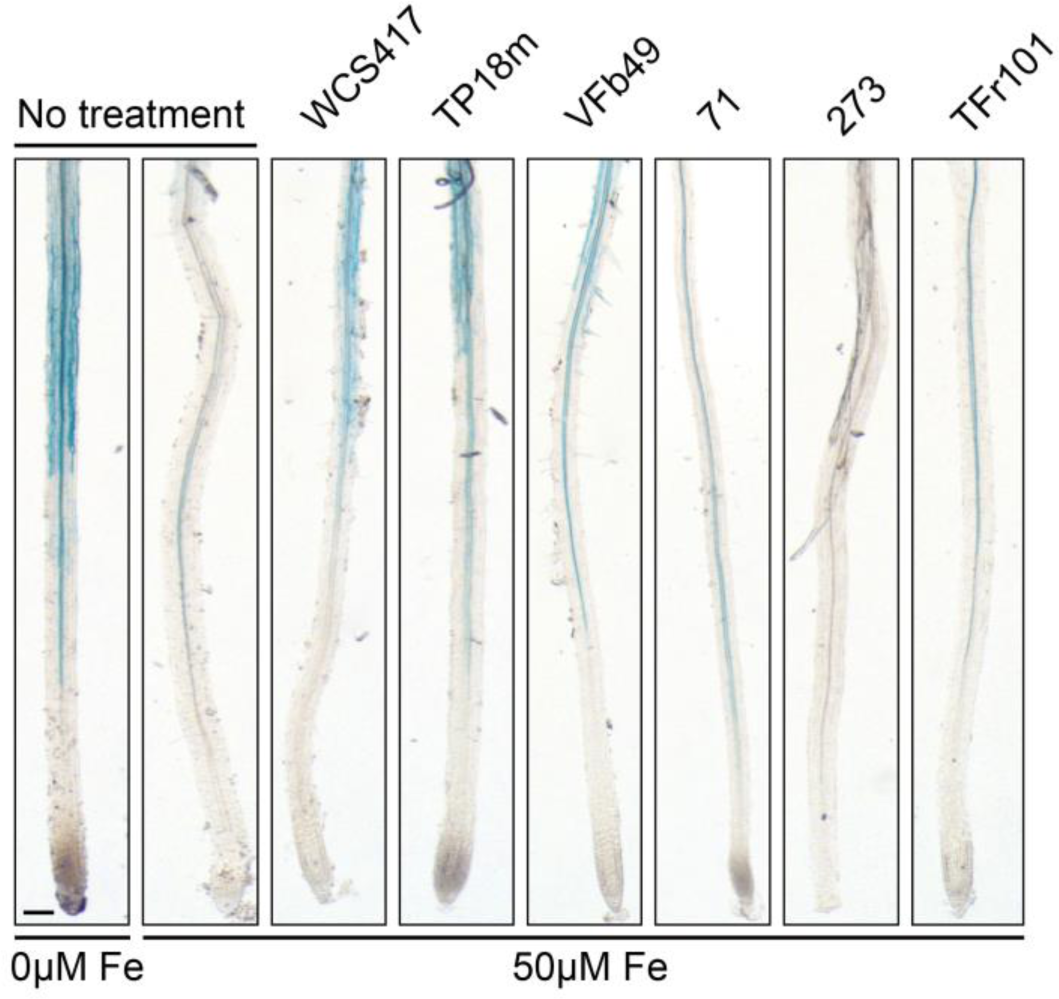
Activation of *MYB72* expression by ISR-inducing bacterial strains. Representative photos of GUS staining of Arabidopsis pMYB72:GFP-GUS plants following bacterial treatments. Media was supplemented with 50 µM FeEDTA (right) to suppress MYB72 induction by low iron (0 µM, left). *P. simiae* WCS417 was used as a positive control. Staining was performed 3 days after treatment. Scale bar = 50 µm.

### Characterization of bacterial plant growth promoting abilities

Besides their immunity-enhancing properties, many rhizobacteria also confer PGP effects such as N_2_-fixation, enhancement of the bioavailability of insoluble minerals, or improved root architecture of host plants (Trivedi et al., 2020). Thus, we further characterized our five strains from a PGP point-of-view. Interestingly, only *P. parafulva* TP18m displayed clear phosphate solubilization activity (Fig. 6A and B), while all strains displayed N_2_-fixation capabilities, and siderophore production. All except *P. peoriae* TFr101 exhibited ACC deaminase activity (Fig. 6B).

**Figure 6.**
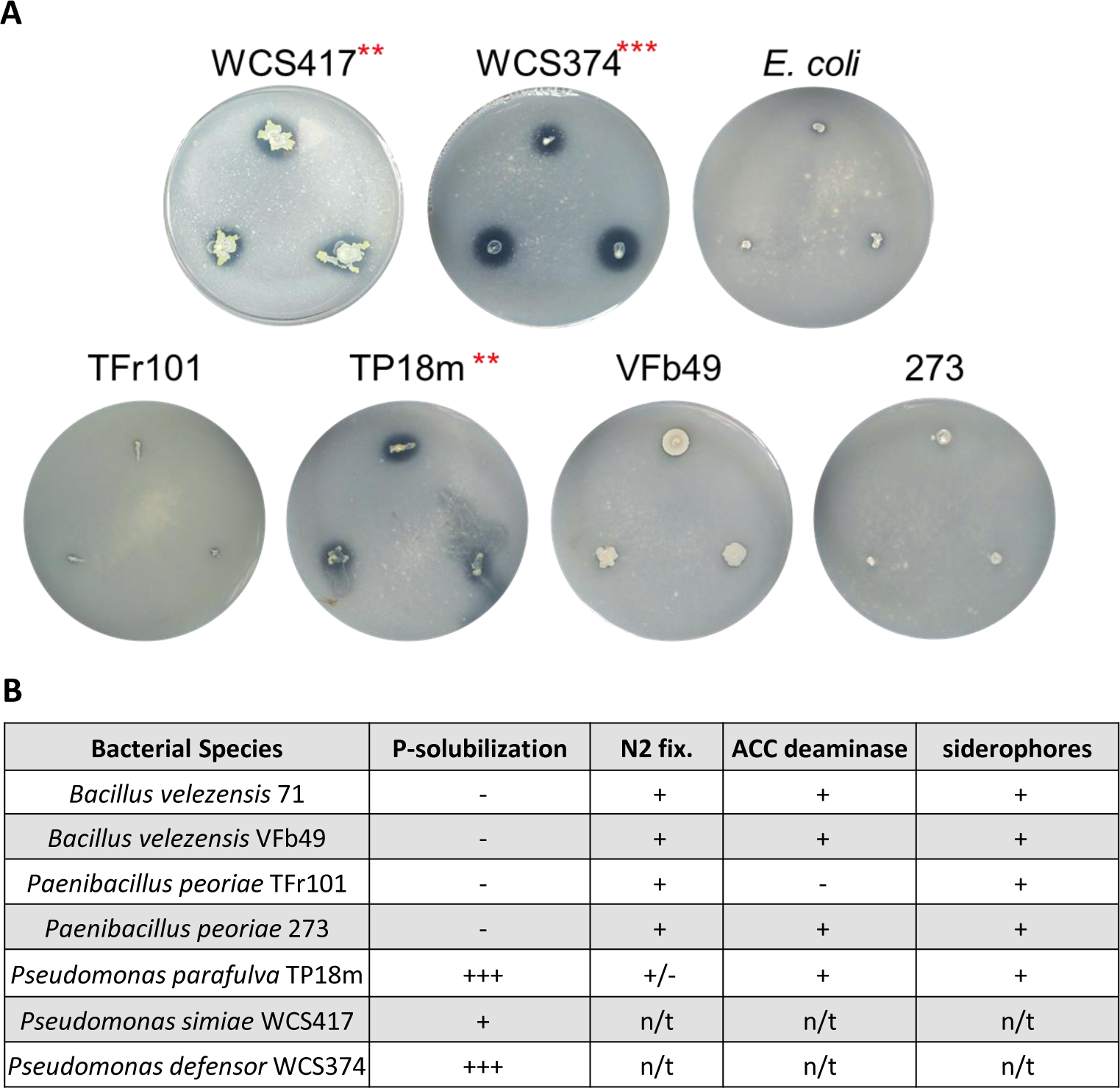
Plant growth promoting effects of bacterial strains. **A.** Phosphate solubilization test on Pikovskaya’s (PVK) agar. Experiments were conducted with 1 plate per strain with 3 biological repeats on each plate. This experiment was conducted 3 times with similar results. Halo formation surrounding the bacteria indicates phosphate solubilization has occurred. **B.** Summary of PGP effects. Measured were Phosphate solubilization, nitrogen fixation, 1-aminocyclopropane-1-carboxylate deaminase (ACC) activity, and siderophore production. n/t, not tested.

These data indicated possible PGP effects by our strains; thus, we conducted *in planta* analysis of the 5 strains to observe effects on root growth and development. Interestingly, only *P. parafulva* TP18m, which also exhibited clear phosphate solubilization activity, exhibited a very strong, and statistically significant effect on root growth and development in both Arabidopsis and tomato (Fig. 7). One of the most common PGPR effects are morphological changes of the root architecture (Zamioudis et al., 2013). Frequently, a stimulation on lateral root formation is observed after treatment with beneficial bacteria. In Arabidopsis, we found that ISR inducing *P. simiae* WCS417 had a mild, but not significant, increase in the lateral root number (LR) compared to the buffer control (Mock) (Fig. 7C). While our non-rhizosphere bacterial control *E. coli* did increase LR formation, *P. parafulva* TP18m had the highest number of LR per plant (Fig. 7A, C).

**Figure 7.**
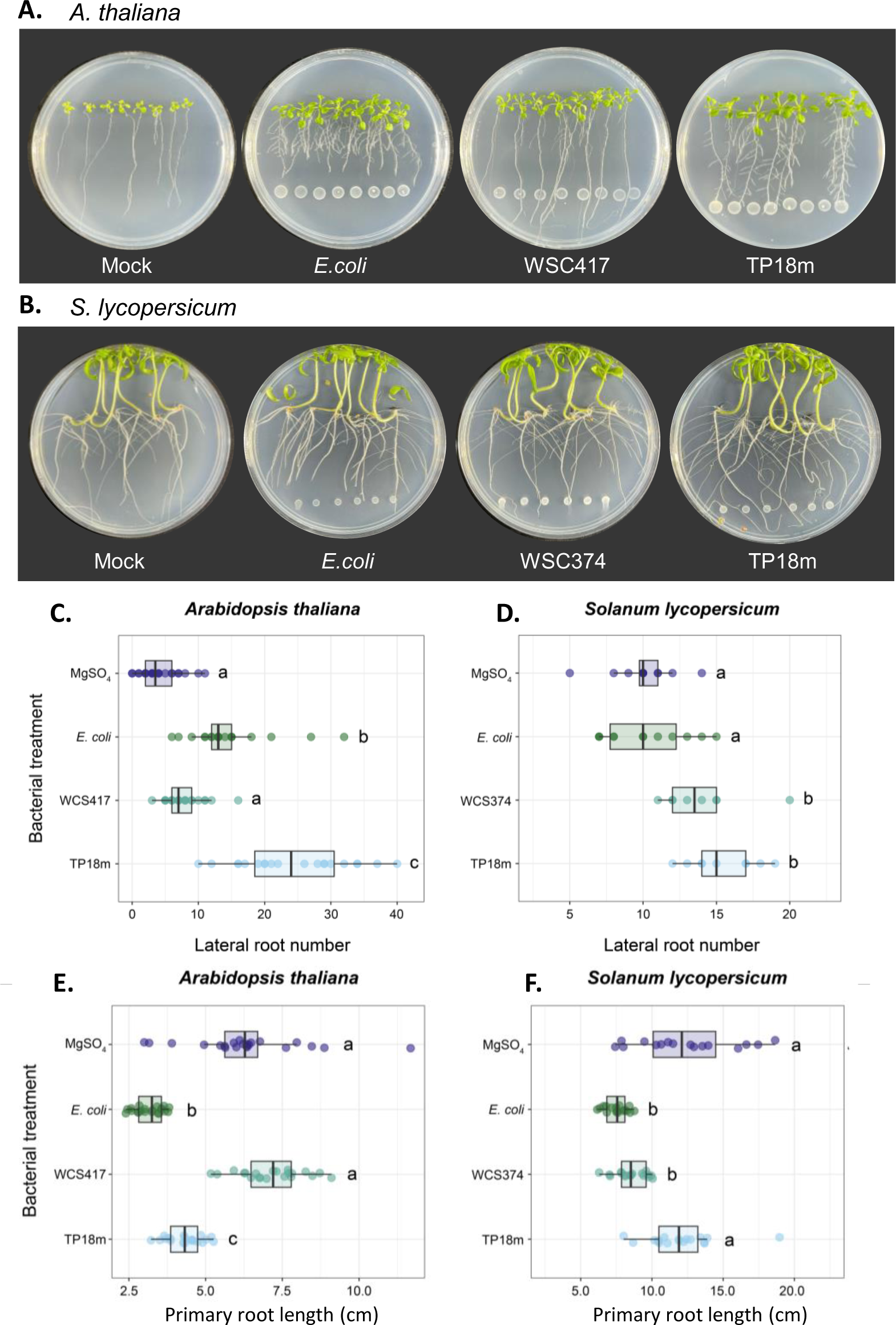
Plant Growth Promoting effects on *Arabidopsis thaliana* and *Solanum lycopersicum.* **(A)** Arabidopsis and **(B)** tomato seedlings were grown on MS agar plates where droplets of MgSO_4_ (mock), *E. coli*, *P. simiae* WSC417 or *P. parafulva* TP18m were spotted at the bottom of the plates. Images were taken at 14 days and **(C, D)** lateral root formation and **(E, F)** primary root length was determined. Differences between MgSO_4_ and each bacterial treatment were evaluated between treatments using a one-way ANOVA followed by a followed by Tukey’s HSD. (n = 6-22; different letters indicate statistically significant differences).

In many cases, plant roots display primary root shortening in response to living bacteria or PAMPs such as flg22 (Gomez-Gomez et al., 1999; Stringlis et al., 2018a) as a part of immune activation. This was clearly the case for *E. coli* (Fig. 7A, E). As previously reported, *P. simiae* WCS417 can actively suppress this effect (Stringlis et al., 2018a) and primary roots exhibited no growth inhibition with *P. simiae* WCS417 (Fig. 7 E). For *P. parafulva* TP18m, although the effect was not as strong as *P. simiae* WCS417, Arabidopsis primary roots were significantly longer than in the presence of *E. coli* (Fig. 7A, E).

In tomato, we observed similar trends. Both *P. defensor* WCS374 and *P. parafulva* TP18m significantly increased the number of LR, while no effect was observed for *E. coli* (Fig. 7D). Primary root growth retardation was seen for *E. coli* and *P. defensor* WCS374 compared to the buffer control but was completely absent for *P. parafulva* TP18m (Fig. 7B, F). Thus, overall, one of our strains, *P. parafulva* TP18m, had a clear PGPR effect on both Arabidopsis and tomato seedlings and suggests an ability to increase overall root biomass.

## Discussion

Plant roots and their surrounding microorganisms in the soil form an intricate relationship where, similarly to the human gut microbiome, beneficial microorganisms promote plant health while pathogenic microorganisms harm the plant (Bakker et al., 2013; Paasch and He, 2021). Therefore, plants can attract beneficial microorganisms to the rhizosphere where they can protect plants either through direct competition with other microbes or via triggering a systemic activation of the plant immune system (Pieterse et al., 2014). To attract beneficial bacteria, plants secrete a wide range of bioactive molecules into the rhizosphere. These include attractants, stimulants and repellants, and the composition varies widely between plant species (Hartmann et al., 2009; Hu et al., 2018). For example, the coumarin scopoletin exhibits selective antimicrobial activity towards soil bacteria and selects beneficial rhizobacteria (Stringlis et al., 2018). Besides these effects, a range of rhizobacteria also can promote plant health by stimulating changes in root architecture, which leads to improved plant performance. These bacteria are termed Plant Growth Promoting Rhizobacteria (Backer et al., 2018).

In the face of an increasing demand due to a growing population and awareness of the potential negative side effects of current industrial agriculture practices, there is increasing interest in more holistic agricultural methods, such as the usage of native soil bacteria to promote plant growth as well as pathogen resistance (Backer et al., 2018; Lahlali et al., 2022).

To this end, we established a method to screen a library of bacteria derived from agricultural soil samples from different geographic regions of Canada (Olishevska et al., 2023) for their ability to induce systemic resistance in tomato plants against the agriculturally important pathogen, *B. cinerea*. Here, we report the detailed characterization of five strains that had been shown to possess antimicrobial properties against *Xanthomonas* species. We found that pretreatment of roots with these bacteria led to decreased *Botrytis*-induced lesion size on leaves. These are two strains of *B. velezensis*, two strains of *P. peoriae*, and one strain of *P. parafulva*. *B. velezensis* belongs to operational group *B. amyloliquefaciens* (OGBa), which contains four *Bacillus* species *(Bacillus amyloliquefaciens, Bacillus siamensis, Bacillus velezensis* and *Bacillus nakamurai*, which exhibit a wide range of plant-beneficial effects (Fan et al., 2018; Ngalimat et al., 2021; Rabbee et al., 2019). They are also closely related to *B. subtilis*, which includes some of the best characterized beneficial rhizosphere bacterial strains (Blake et al., 2021). *Paenibacillus* species are increasingly associated with plant growth promotion (Wang et al., 2023) as well as biocontrol capabilities against e.g., *Coniella vitis* (white rot of grape) and *Fusarium* spp. (Yuan et al., 2022), among others. Like strain TP18m, other *P. parafulva* strains also display antimicrobial activities (Abanda-Nkpwatt et al., 2006; Kakembo and Lee, 2019).

Beneficial rhizobacteria can protect plants in more than one way: One is through competition with other bacteria in the rhizosphere (via antimicrobial activity), another is by activating the plant immune system both in the root as well as systemically in aboveground tissues (Pieterse et al., 2012; Trivedi et al., 2020), a phenomenon termed Induced Systemic Resistance (ISR). ISR triggered by several *Pseudomonas* species has been well-characterized and involves JA and ET pathways (Berendsen et al., 2015; Pieterse et al., 2014). However, it is not clear whether other bacterial species also act through the same pathway; there are some reports suggesting alternative pathways (Nie et al., 2017; Niu et al., 2011; van de Mortel et al., 2012; Vlot et al., 2021). Thus, we characterized our five strains from several angles and found diverse capabilities in terms of antimicrobial activities and their effects on plants.

To gain an insight in a mechanistic aspect, we took advantage of various Arabidopsis mutant lines. Interestingly, only pre-treatment with *B. velezensis* VFb49 and *P. parafulva* TP18m reduced the *Botrytis* lesion size in wildtype Arabidopsis (Fig. 4). Accordingly, only these two strains induced the expression of the ISR marker *MYB72* in Arabidopsis (Fig. 5), suggesting that only these two are triggering the canonical ISR pathway in the same way as *P. simiae* WCS417 does (Pieterse et al., 2021). However, the other three strains clearly induce systemic immunity in tomato. This suggests that a compatibility between specific bacterial and plant host species is required (Berendsen et al., 2015). *P. defensor* WCS374 was effective in tomato. On the other hand, *P. defensor* WCS374 is not a strong inducer of ISR in Arabidopsis (Berendsen et al., 2015). Different affinities of WCS374, WCS358 and WCS417 to induce ISR were also reported in radish and tobacco (Leeman 1996; Van Loon et al., 2008).

Which factors determine this host specificity is currently unknown; however, since we tested tomatoes and Arabidopsis under the same conditions, it is certainly related to the host plant species. Furthermore, ISR establishment may require a high degree of compatibility. For example, WCS417 triggers ISR in the Arabidopsis Col-0 and Ler-0 accessions but fails to do so in Ws-0 and RLD1 (Ton et al., 2001; Van Wees et al., 1997), suggesting genetic compatibility, even within the same species, is an important factor. Some bacteria trigger ISR in multiple species, while others fail to do so (Berendsen et al., 2015; Salwan et al., 2023), so the efficacy of each strain has to be evaluated for each crop.

Recent data clearly shows that beneficial rhizobacteria also trigger a PTI response, but many bacteria can suppress the downstream immune response; at the same time, both the induction and dampening of the immune response is required for successful colonization (Ma et al., 2021; Millet et al., 2010; Stringlis et al., 2018a; Tzipilevich et al., 2021; Yu et al., 2019). The production of ROS is a hallmark of PTI responses (DeFalco and Zipfel, 2021) and is important for root colonization of *B. velezensis* FZB42 and auxin secretion of the host plant (Tzipilevich et al., 2021). Song et al. (2021) recently proposed that ROS production mediated by the receptor kinase FERONIA controls root colonization by Pseudomonads. We also found that our strains failed to induce ISR in the ROS mutant *rbohD*, suggesting the importance of ROS production for ISR establishment as well. The two ISR-strains also were dependent on JA/ET signaling, rather than SA, as reported for *P. simiae* WCS417 (Pieterse et al., 2021), suggesting they trigger the canonical ISR pathway.

The nature of the signalling molecule(s) that, following ISR induction in the roots, moves to the aboveground tissues of the plant remains unclear. Recently, lipid transfer proteins like EARLI1 or AZI1, which are well-studied components of SAR long distance signaling (Vlot et al., 2021), have been connected to ISR (Cecchini et al., 2015). DIR1 forms a complex with AZI1 and EARLI1 (Cecchini et al., 2015; Yu et al., 2013). Thus, we tested the SAR-deficient *dir1* mutant (Maldonado et al., 2002). Again, all three strains could not trigger ISR in *dir1*, supporting the notion that there is some overlap between SAR and ISR signaling. AZI1 and EARLI1 carry the SAR-related compound azelaic acid (AZA) to systemic tissues. So, potentially a lipid signal could also play a role in ISR. The involvement of lipids in rhizobacteria-root interaction is just starting to be investigated (Macabuhay et al., 2022).

While only *P. parafulva* TP18m led to statistically significant root architecture changes in both tomato and Arabidopsis, we also observed some trends for *B. velezensis* VFb49 and *P. peoriae* TFr101. As previously reported, *P. simiae* WCS417 suppressed the PTI response against bacteria/PAMPS (root shortening) and allowed the root to grow through the bacterial culture (Stringlis, et al., 2018), while we observed no strong effect on LR formation. On the other hand, *P. parafulva* TP18m induced a dramatic root architecture change with somewhat shorter but highly branched lateral roots. This resembles results by Li et al. (2022), who also observed longer, but less branched roots in the presence of *P. simiae* WCS417 and a shorter but highly branched root phenotype in the presence of *Pseudomonas Castanea mollissima* (strain 11). Thus, different PGPRs exert different effects on their host plants and the combination of strains (i.e. generation of consortia) with complementary effects can boost plant performance even further (Li et al., 2022).

In conclusion, we established a screening method and characterized five of the identified strains in detail. Further characterization revealed that these strains possess diverse characteristics in terms of their properties and modes of action. Only two also triggered the canonical ISR pathway in Arabidopsis and one exhibited strong PGPR effects. Future work is needed to elucidate whether the other strains are tomato-specific or have a wider host range. The diverse characteristics of these strains can be a valuable tool to analyse the different signaling pathways that these strains activate in plants, both with respect to immunity and PGP effects. Furthermore, the characterization of these strains – and other strains that will be discovered through our screen – will be a foundation to assemble bacterial consortia that will hopefully be able to provide robust protection for crops in the field.

## Supplemental Data

**Table S1. Strain-specific primers used in this study.**

## Acknowledgements

We would like to thank Cara Haney (University of Pittsburgh) for the MYB72:GUS seeds, Robin Cameron (McMaster University) for *dir1* seeds, and UAMH Center for Global Microfungal Biodiversity and Agriculture Canada for providing and maintaining our Botrytis strain.

## Author Contributions

WM, ED and KY conceived the research. WM, TB, ED, and KY designed the research. MEWL, WY, NML and MT conducted research at UofT, MCG and AN conducted the experiments at INRA. MEWL and WY analyzed data. WM, MEWL and KY wrote the article.

## Conflicts of Interest

The authors declare that they have no competing interests.

## Funding

This work was supported by an NSERC Strategic Partnership Grant to KY and ED, and an Ontario Graduate Scholarship to MEWL.

## Abbreviations

ISR: Induced Systemic Resistance
SAR: Systemic Acquired resistance
DIR1: defective in induced resistance1
SA: salicylic acid
JA: Jasmonic acid
ET: ethylene
PGP(R): plant growth promoting (rhizobacteria)
EARLI1: EARLY ARABIDOPSIS ALUMINUM INDUCED 1
AZI1: AZELAIC ACID INDUCED 1
PAMP: pathogen associated Molecular pattern
PTI: PAMP-triggered immunity
ROS: Reactive Oxygen Species

## Notes

### Competing Interest Statement

The authors have declared no competing interest.

